# Estimating plasmid conjugation rates: a new computational tool and a critical comparison of methods

**DOI:** 10.1101/2020.03.09.980862

**Authors:** Jana S. Huisman, Fabienne Benz, Sarah J.N. Duxbury, J. Arjan G.M. de Visser, Alex R. Hall, Egil A.J. Fischer, Sebastian Bonhoeffer

## Abstract

Plasmids are important vectors for the spread of genes among diverse populations of bacteria. However, there is no standard method to determine the rate at which they spread horizontally via conjugation. Here, we compare commonly used methods on simulated and experimental data, and show that the resulting conjugation rate estimates often depend strongly on the time of measurement, the initial population densities, or the initial ratio of donor to recipient populations. Differences in growth rate, e.g. induced by sub-lethal antibiotic concentrations or temperature, can also significantly bias conjugation rate estimates. We derive a new ‘end-point’ measure to estimate conjugation rates, which extends the well-known Simonsen method to include the effects of differences in population growth and conjugation rates from donors and transconjugants. We further derive analytical expressions for the parameter range in which these approximations remain valid. We present an easy to use R package and web interface which implement both new and previously existing methods to estimate conjugation rates. The result is a set of tools and guidelines for accurate and comparable measurement of plasmid conjugation rates.

**Highlights:**

- We present an R package and Shiny app to estimate conjugation rates from experimental data
- We highlight the diversity of existing measures used to estimate conjugation rates
- We show these measures are biased for populations with differing growth and conjugation rates from donors or transconjugants
- We develop a new ‘end-point’ measure which accounts for such differences in growth and conjugation rates

## 1 Introduction

Plasmids are extra-chromosomal, self-replicating genetic elements that can spread between bacteria via conjugation. They spread genes within and between bacterial species and are a primary source of genetic innovation in the prokaryotic realm [1, 2]. Genes disseminated by plasmids include virulence factors, heavy metal and antibiotic resistance, metabolic genes, as well as genes involved in cooperation and spite [2, 3, 4, 5]. To understand how these traits shape the ecology and evolution of bacteria [6], it is of fundamental importance to understand and quantify how plasmids spread.

The maintenance and spread of a plasmid in a population is determined by two factors: (i) the horizontal transmission of plasmids between neighbouring bacteria (conjugation) and (ii) the vertical transmission of a plasmid with its host upon cell division (clonal expansion). Plasmid conjugation requires physical contact between donor cells (D), carrying the plasmid, and recipient cells (R), to create transconjugant cells (T), i.e. recipients carrying the plasmid [1]. The transconjugants then further contribute to the transfer of the plasmid to recipients. The conjugation rates from transconjugants can be substantially higher due to transitory derepression of the conjugative pilus synthesis [7, 8], and because transconjugant and recipient cells are the same strain with the same restriction modification systems [9, 10]. In addition, the rates of clonal expansion of D, R, T populations can differ strongly, especially when the plasmid is transferred across species boundaries [9].

Quantifying the horizontal and vertical modes of plasmid transmission separately is important for the fundamental understanding of plasmid biology and plasmid-host interactions, as well as the prediction of plasmid spread and selection in diverse environments. The burden of a plasmid on its host cell, the selection of plasmid-borne traits, and the regulation of plasmid conjugation machinery may all be affected by mutations and (a)biotic factors in the environment. Their interdependence can be assessed only when growth and conjugation are quantified independently. This is of particular importance for the epidemiology of antibiotic resistance plasmids for instance, where plasmids with intrinsically high conjugation rates necessitate different interventions than clonally spreading plasmid-strain associations [11, 12].

Given the importance of plasmid spread, it is surprising that there is no generally accepted method to quantify the amount of conjugation that occurs between bacterial populations. Differences between conjugation assays are dictated by the variety of biological systems in which conjugation occurs: different species require different growth medium for instance, and some plasmids require solid matrices for conjugation. All conjugation assays have in common that the donor and recipient cells are cultured together in or on a specific growth medium for a certain amount of time *t*. After this time, the resulting population densities are measured. However, assays differ in which populations are measured - D, R, and T, or only a subset thereof [9, 13, 14]; in the experimental system used - e.g. well-mixed liquid cultures, filters, plates, the gut of vertebrate hosts [9, 15, 16]; the duration of the assay *t* - from 1 hour to multiple days [17, 18]; and the way population densities are measured - e.g. through selective plating, or flow cytometry [15, 19, 20]. Differences in the output of such conjugation assays are then further exacerbated when the measured population densities are related to the amount of conjugation that occurred. Indeed, there is no consensus on what to call this quantity: commonly used phrases include conjugation frequency [21, 22], plasmid transfer rate constant [13, 23], or transfer efficiency [19, 20]. More than 10 different methods to quantify conjugation are currently found in the literature (see Table 1).

**Table 1:**
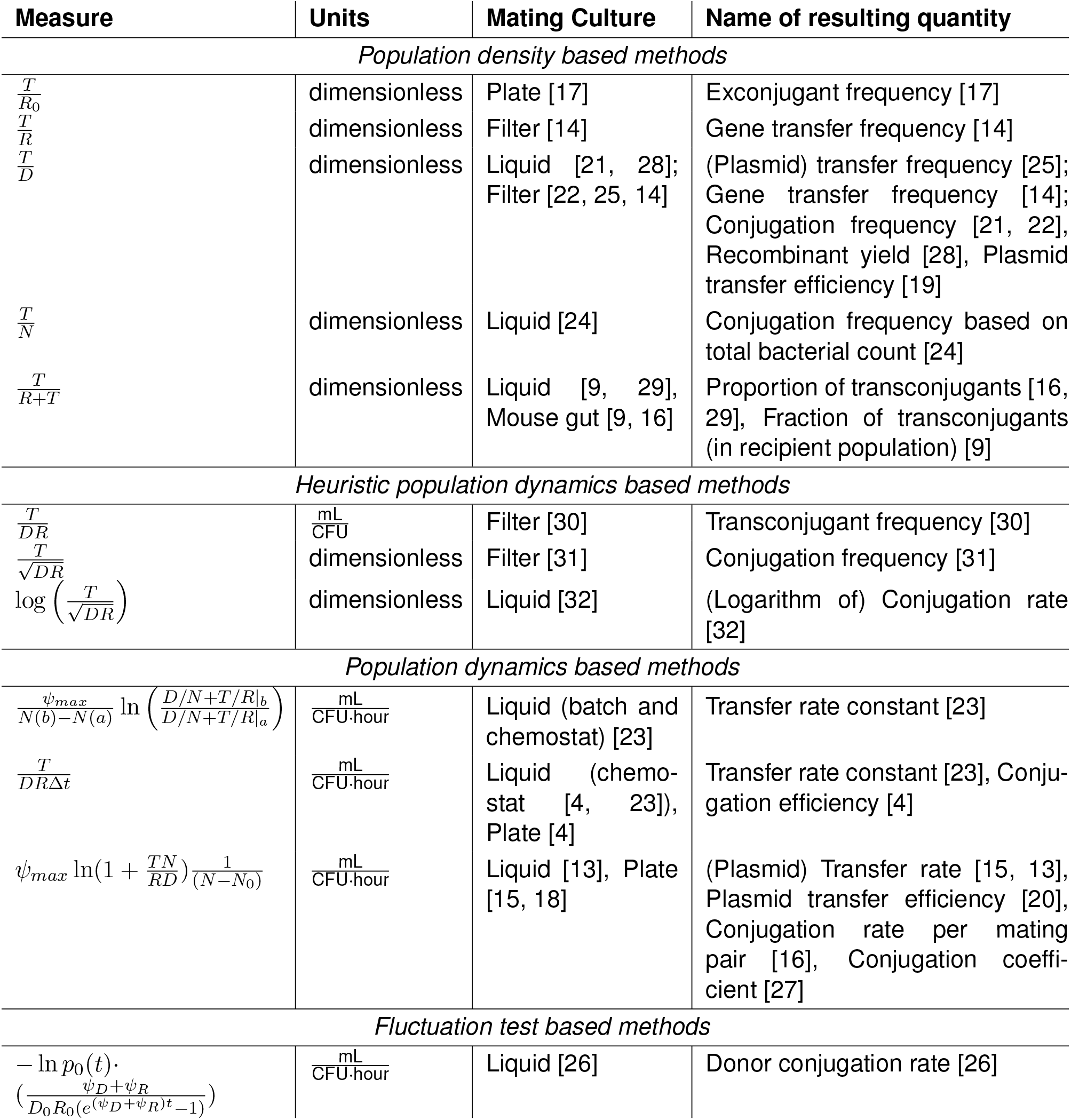
Measures of conjugation proficiency reported in the literature. Here *D, R, T* stands for the population density of donors, recipients, and transconjugants at the time point of measurement, *N* is the total population density (*N* = *D* + *R* + *T*), *N*_0_ is the initial total population density, *R*_0_ is the initial population density of recipients, *ψ_max_* is the maximum growth rate of the mating culture, and *p*_0_(*t*) is the probability that zero transconjugants are observed at time *t*.

Many methods are based on the ratio between population densities, such as *T/D* or *T/R*, to quantify the fraction of transconjugants at the end of the conjugation assay (these will collectively be referred to as ‘population density based methods’) [8, 17]. However, these measures vary as a function of the initial population densities, the initial donor to recipient ratio, and the length of the conjugation assay [13, 18]. In addition, they are affected both by a plasmid’s transfer rate, as well as its clonal expansion [19]. Thus, experimental results reported with such measures are not comparable between studies without detailed information on the experimental conditions and growth rates of the strains involved (which are often lacking) [15, 18]. The resulting measurements are also not a *priori* comparable across experimental conditions that could affect the growth rate, including differing nutrient conditions [24], recipient species [9, 17, 25], temperatures [21], and (sub-lethal) antibiotic exposure [22]. This limits the predictive power of conjugation proficiency when expressed as a ratio of population densities [19].

Population dynamic models were developed specifically to disentangle the influence of horizontal and vertical plasmid transmission on final population density. In 1979, Levin et al. showed that conjugation in well-mixed liquid cultures can be accurately described with the mass action kinetics also used to describe chemical reactions [23]. They described a method to estimate the conjugation rate from bacterial population densities using linear regression in the exponential or stationary growth phase [23]. This method was developed further by Simonsen et al. [13], who derived a closed formula for the conjugation rate. They call this an ‘end-point’ method since it requires a single measurement of *D, R* and *T* population densities at the end of the conjugation assay, as opposed to time-course data. The method further requires knowledge of the mating population growth rate and initial population density. Recently a new method was introduced which explicitly takes into account the stochasticity of conjugation [26]. This so-called ‘low-density method’ still assumes mass action kinetics of plasmid transfer, as well as deterministic and exponential growth of the donor and recipient populations. However, the stochastic treatment has the advantage of allowing shorter incubation times and shows low variance of the estimate across experimental replicates.

Although the Simonsen method is widely regarded as the most robust method available to estimate conjugation rates [18], more than thirty years after its publication an astounding variety of methods is still in common use (see Table 1). One can speculate whether this slow adoption of the Simonsen method has been because of a sense of unease with the model-based formulation, the extra bit of work involved in measuring the population growth rate, or the power of habit in using population density based methods. In addition, the Simonsen method has the drawback that it does not account for differences in growth rates between strains, nor in differences in conjugation stemming from donors or transconjugants. Fischer et al. [27] extended the Simonsen model along these lines, but their approach requires time course measurements and a fitting procedure which is sensitive to the initial values of the optimisation. Thus, there is a clear need to reiterate the drawbacks of population density based methods, and to lower the barrier to widespread use of better population dynamics based alternatives.

Here, we show the limitations of existing measures of conjugation proficiency on simulated and experimental data, including their dependence on measurement time point, as well as the initial population densities and ratios. To mitigate these limitations, we extend the Simonsen model to include the effects of differential population growth and conjugation rates from donors and transconjugants. For this extended model we derive a new formula for the conjugation rate, as well as the critical time within which these approximations are valid. We show how our extended model compares to the original Simonsen model as a function of differences in the growth and conjugation rates. We further developed an R package and web interface (a Shiny app) to facilitate the calculation of a variety of conjugation rate methods from experimental data and to allow testing whether these were measured within the critical time. The result is a set of guidelines for easy, accurate, and comparable measurement of conjugation rates and tools to verify these rates.

## 2 Materials and Methods

### Simulations

We simulated bacterial population dynamics under the extended Simonsen model (ESM, described in the Theory and Calculations section) and evaluated the performance of different conjugation rate estimation methods. The code to simulate the different models, as well as the specific parameter settings for each figure are available from https://github.com/JSHuisman/conjugator_paper.

### Conjugation Experiments

#### Strains

We used *Escherichia coli* strains for the time course and full protocol conjugation assays. To distinguish the conjugation rate from donors (*γ_D_*) from that of transconjugants (*γ_T_*), the full protocol (described in the Results section) consists of two experiments: (i) the DRT assay combines donors (D) and recipients (R) to form transconjugants (T), and (ii) the TRT assay combines labelled transconjugants with recipients (both of the same genetic background) to form 2nd generation transconjugants. We recently described the E. *coli* strains used in these conjugation assays but provide details of their identity here [33]. For DRT assays, the Donor (D) was a natural chicken isolate (ESBL-375) carrying an IncI1 plasmid with the cefotaxime resistance gene *bla_CTX-M-1_* (gift from Michael Brouwer, Wageningen Bioveterinary Research, The Netherlands). As Recipient (R) we used strain DA28100: a MG1655 laboratory strain chromosomally labelled with chloramphenicol resistance in the *galK* locus (MG1655, *galK::sYFP2opt-cat*; GenBank accession number KM018300) [34]. Strain DA28100 was a kind gift from the Dan Andersson lab via Peter A Lind, constructed by Erik Gullberg/Wistrand-Yuen. For TRT assays, we derived the Donor (D’) strain from strain DA28100 by removing the *cat* gene flanked by FLP recombinase target (FRT) sites, by expression of FLP recombinase (strain constructed by Andrew Farr, Arjan de Visser lab). This strain was isogenic to strain DA28200 described in Gullberg *et al*. [34], constructed via the same method. To distinguish strain D’ from strain R, a spontaneous nalidixic acid (NAL) resistant mutant of strain D’ was isolated. Strains R and D’ were otherwise isogenic. Strain D’ received the IncI1 plasmid from ESBL-375 via a prior conjugation assay.

Compared to the full protocol described in the main text, we decided to label strain D’ (rather than R) for use in the TRT assay, since this allowed use of the same recipient strain (R) in both DRT and TRT assays and the simultaneous measurement of conjugation and growth rates of all strains in the same assay. The TRT assay is thus more correctly labelled the D’RT assay.

Strains were cryopreserved by storing in 20% v/v glycerol in LB medium (10 g L^−1^ (bacto) tryptone, 5 g L^−1^ yeast extract, 10 g L^−1^ NaCl) at −80°C [33]. Strains were revived prior to growth or conjugation assays by streaking to single colonies on agar plates (LB or VL medium as described in assays below). Due to cryopreservation and revival, any transient de-repression state in strain D’ has likely been lost.

#### Time course conjugation assay

To compare the sensitivity of different conjugation rate estimation methods to the measurement time point, we performed a DRT conjugation assay with sampling at multiple time points. Overnight cultures were incubated at 37°C in LB medium with shaking at 250 rpm. Cultures were then diluted in LB medium and grown into early stationary phase. Strains D and R were mixed in a 50:50 ratio and 100 *μ*L of culture was diluted into 10 mL LB medium in a 50 mL falcon tube, resulting in an approximate initial density of 1.5 · 10^7^ CFU/mL per strain. The culture was vortex-mixed and sampled for plating on selective agar at *t* = 0. The culture was then incubated under static conditions at 37°C, vortex-mixed and sampled after 0.75, 1.5, 5, 19 and 25 hours during the conjugation assay. We chose to use static rather than shaken conditions to maximize the number of successful conjugation events, as D-R mating pairs may be broken with agitation [35]. Only after 19 and 25 hours we observed some cell settlement prior to mixing, which may have limited numbers of cell-cell contacts formed. Following serial dilutions, CTX plates (cefotaxime, 1 mg/L) were used to select for strain D and transconjugants, CAM (chloramphenicol, 32 *μ*g/mL) for strain R and transconjugants and CTX+CAM for transconjugants only. Cell densities (CFU/mL) were calculated based on the dilution factors and transconjugant counts were subtracted from total counts on the CTX and CAM plates. We assume that the transconjugant counts on double selective plates reflect the formation of transconjugants in liquid culture, however we did not control against the possibility of ‘on-agar’ conjugation events on the double-selective agar plates. Such ‘on-agar’ conjugation could have increased transconjugant counts [9, 36, 37], but would have done so to an equal extent across all plated time points. We verified that the CTX and CAM agar types were selective for strain D and strain R respectively by plating a monoculture of each strain on the two agar types. We did not see any colony growth on the opposite selective agar for each strain. The mixed culture experiment was repeated in three biological replicates (separate experimental runs) with three agar plate replicates per time point where possible. Cell densities across replicate agar plates were averaged for each biological replicate per time point.

We then performed a growth assay of strains D and R, and three transconjugant clones (T). A single transconjugant clone from each of the three replicate conjugation assays was selected and strains were grown overnight in LB medium. Three biological replicate cultures of strains D and R were prepared. The growth assay was set up in a 96-well microtiter plate. Note that these growth conditions (96-well microtiter plate) differ from the conjugation assay described (culture tubes) and therefore might have caused the growth rates to differ in the conjugation assay. However, because the growth rates of D and R strains were both measured in microtiter plates, we do not expect large effects on fitness cost and conjugation rate estimates. From overnight cultures, a 1:100 dilution was performed by adding 2 *μ*L cells in 200 *μ*L of LB medium in each of six (technical) replicate wells within a column of the 96-well microtiter plate. This resulted in initial cell densities of approximately 10^7^ CFU/mL. OD_600_ readings were measured in a Victor3 plate reader (Perkin and Elmer, Massachusetts, US) every 16 minutes over 24 hours during incubation at 37°C with orbital shaking prior to each reading. Growth rates were estimated per strain, per biological replicate by pooling data from each set of six replicates. OD data was clipped after 8 hours (when strains entered stationary phase) and a logistic growth model without lag phase was fitted to each set of data as described in [27] using non-linear least squares fitting in R version 4.0.3.

#### Full protocol

To illustrate the full protocol, we ran DRT and TRT conjugation assays with strains D and R, and D’ and R respectively. Conjugation and growth assays were run in a single 96-well microtiter plate, including growth profiling of the mixed conjugation cultures (N). To measure growth of a transconjugant strain (T), a transconjugant was isolated from a prior conjugation assay in which the IncI1 plasmid was transferred from strain D to strain R. The assays were repeated in three biological replicates (separate days). The methods described below for the growth and conjugation assays are similar to those described by Duxbury *et al*. [33]. The TRT (D’RT) assay performed here matches one of the control assays performed by Duxbury *et al*. [33] (1.0x_VL medium condition) in which monoculture growth rates were measured alongside conjugation rates after 4 hours.

Overnight cultures were prepared in VL (Viande-Levure) medium [38]. A separate culture per strain was then diluted by 1:100 and grown into exponential phase at 37°C and 250 rpm until OD_600_ nm (measured via a spectrophotometer) reached 0.4 (approximately 1.5 hours). Each strain was then diluted to 2 · 10^6^ CFU/mL in VL medium (approximately 100-fold dilution). monocultures and mixed cultures were prepared with the same starting density per strain. For each monoculture, the 2 · 10^6^ CFU/mL culture was diluted two-fold and 200 *μ*L aliquots were added in two replicate microtiter wells. For mixed cultures, strain cultures (2 · 10^6^ CFU/mL) were mixed in 50:50 ratios and a single 200 *μ*l aliquot of each was added to the microtiter plate. OD_600_ readings were measured in the Victor3 plate reader at 37°C under static conditions every 6 minutes up to 24 hours. OD_600_ data was blank corrected with the minimum OD reading of a medium control well. Maximum growth rates were estimated from the best-fit maximum gradient of natural logarithm-transformed OD values during sliding 1.5-hour windows in the exponential phase (first four hours of growth), similarly to the methods of [39], using Python version 3.6.3. To enumerate strain densities, the mixed cultures were serially diluted appropriately and plated on selective VL agar at *t* = 0, 4 and 24 hours. Two agar plate replicates were included for each agar type per strain, per time point. Cell densities across replicate agar plates were averaged for each biological replicate per time point. Strain D was selected on CTX and strain D’ was selected on CTX+NAL (cefotaxime, 1 mg/L + nalidixic acid, 20 *μ*g/mL). Strain R was selected on CAM and transconjugants were selected on CTX+CAM. Transconjugant counts were subtracted from counts on CTX and CAM plates. Plates were incubated at 37°C for 24 hours. Selectivity of the agar types was confirmed by plating of monocultures at the start and end point of the assay.

#### Data availability

Raw and processed datasets are available via the github repository at https://github.com/JSHuisman/conjugator_paper/blob/master/data/.

## 3 Theory and Calculations

### The Simonsen Model (SM)

Simonsen et al. [13] developed a model (called the SM in the following) that estimates the conjugation rate from end-point measurements of the population densities (*D, R, T*), the initial population density (*N*_0_ = *D*_0_ + *R*_0_), as well as the joint growth rate (*ψ_max_*) of these populations. This model accounts for resource competition between the populations, and the elegant mathematical solution critically requires the assumption that both growth and conjugation have the same functional dependency on the resource concentration. The SM implicitly assumes that conjugation does not occur during the stationary phase. The dynamical equations are given by:

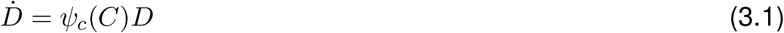

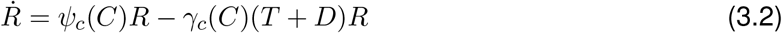

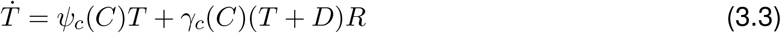

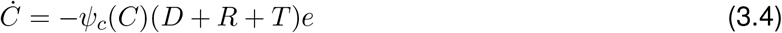

where the designations *D, R, T* stand for donors, recipients, and transconjugants respectively, 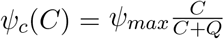 is the growth rate, 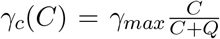 is the conjugation rate, *C* is the resource, and *e* is the conversion factor of resource into cells.

From this model, Simonsen et al. [13] derived that at any time point during the experiment the following relation holds:

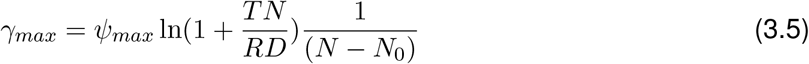

where *N* = *D* + *R* + *T* is the total population density at the measurement time point, *N*_0_ is the initial population density, and the growth rate *ψ_max_* should be determined from the conjugating population during the phase of exponential population growth.

### The Extended Simonsen Model (ESM)

The SM makes two implicit simplifying assumptions. First, it assumes that donors, recipients and transconjugants all have the same growth rate. Second it assumes that the conjugation rate from donors to recipients (*γ_D_*) and from transconjugants to recipients (*γ_T_*) is the same. Both of these assumptions may not be justified. We thus extend the SM to reflect population specific growth rates (*ψ_D_, ψ_R_, ψ_T_*) and conjugation rates (*γ_D_, γ_T_*). This models is called the extended Simonsen model (ESM), and its dynamical equations are:

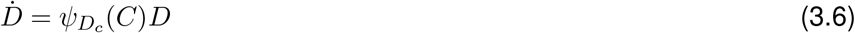

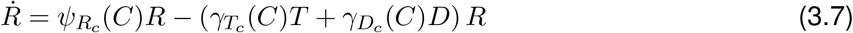

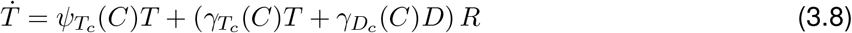

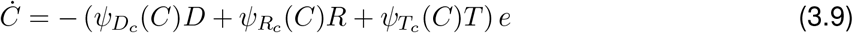

where 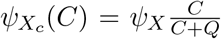 are the population specific growth rates (subscript *X* stands for *D, R, T*), and 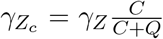 are the conjugation rates from donors or transconjugants (subscript *Z* stands for *D, T*).

### The Approximate Extended Simonsen Model (ASM)

We can simplify the equations for the ESM (eqs. 3.6-3.9) by assuming that the growth and conjugation rates are constant until the resource *C* is gone and switch to zero in stationary phase. This assumption allows one to drop the equation for the resource *C* as long as the stationary phase has not yet been reached. The dynamical equations of the Approximate Extended Simonsen Model (ASM) then become:

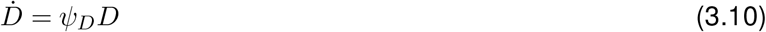

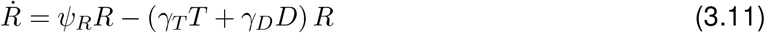

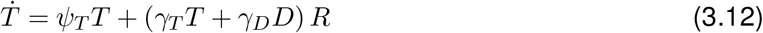

Assuming that initially the dynamics of the recipient population are dominated by growth, i.e. *ψ_R_R* >> *γ_T_TR* + *γ_D_DR*, and that the transconjugant population is not yet dominated by conjugation from transconjugants, i.e. *ψ_T_T* + *γ_D_DR* >> *γ_T_TR*, we obtain that the conjugation rate *γ_D_* at a time point *t* is given by (see Supplementary Materials for detailed derivation):

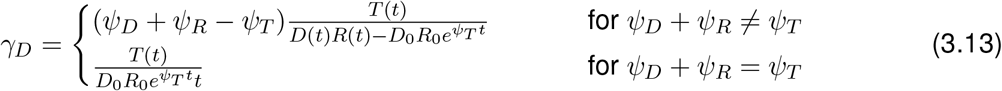

This estimate of the conjugation rate based on the ASM can be used instead of the Simonsen endpoint formula (eq. 3.5) when the growth rates and conjugation rates differ between populations. It is valid as long as the approximate solutions are good approximations to the full ODE. In the Supplementary Materials we derive the critical times (eqs. 9.24, 9.25, 9.26) beyond which different aspects of the approximation are not sufficient anymore, and the ASM formula starts to break down. With ‘the’ critical time *t_crit_*, we refer to the minimum *t_crit_* = min(*t*_*c*1_, *t*_*c*2_, *t*_*c*3_) of these three time points.

## 4 Results

### Population based methods depend sensitively on the experimental conditions

To study the merits of different measures used to quantify conjugation, we test the behaviour of the most common measures on simulated bacterial population dynamics. To this end, we simulate the population dynamics using the extended Simonsen model with resource dynamics (ESM) to include a maximum of biologically relevant detail (see Figure 1A; more scenarios can be investigated on the Shiny app). The population density based measures vary over many orders of magnitude, depending on when the population densities are measured (Figure 1B). Given the simulated cost of plasmid carriage, the *T/D* estimate is higher than *T/*(*R* + *T*), although both would give (approximately) the same result if the growth rate of the *D* and *R* populations were the same. The measure 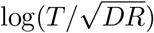 is relatively stable as a function of the measurement time. However, it is negative as long as *T* is smaller than *D* and *R*, and one has to take the absolute value to allow comparison with the other conjugation measures. The measure *T/DR* performs almost as well as the populations dynamics based measures (SM / ASM), as it approximates the same mass action kinetics for short time frames. One can also see that the dimensionless population density based measures are many orders of magnitude larger than conjugation rates estimated using population dynamic models, as the latter are typically reported in mL · CFU^−1^h^−1^. Similar biases are found when applying the same methods to experimental data (Figure 1C).

**Figure 1:**
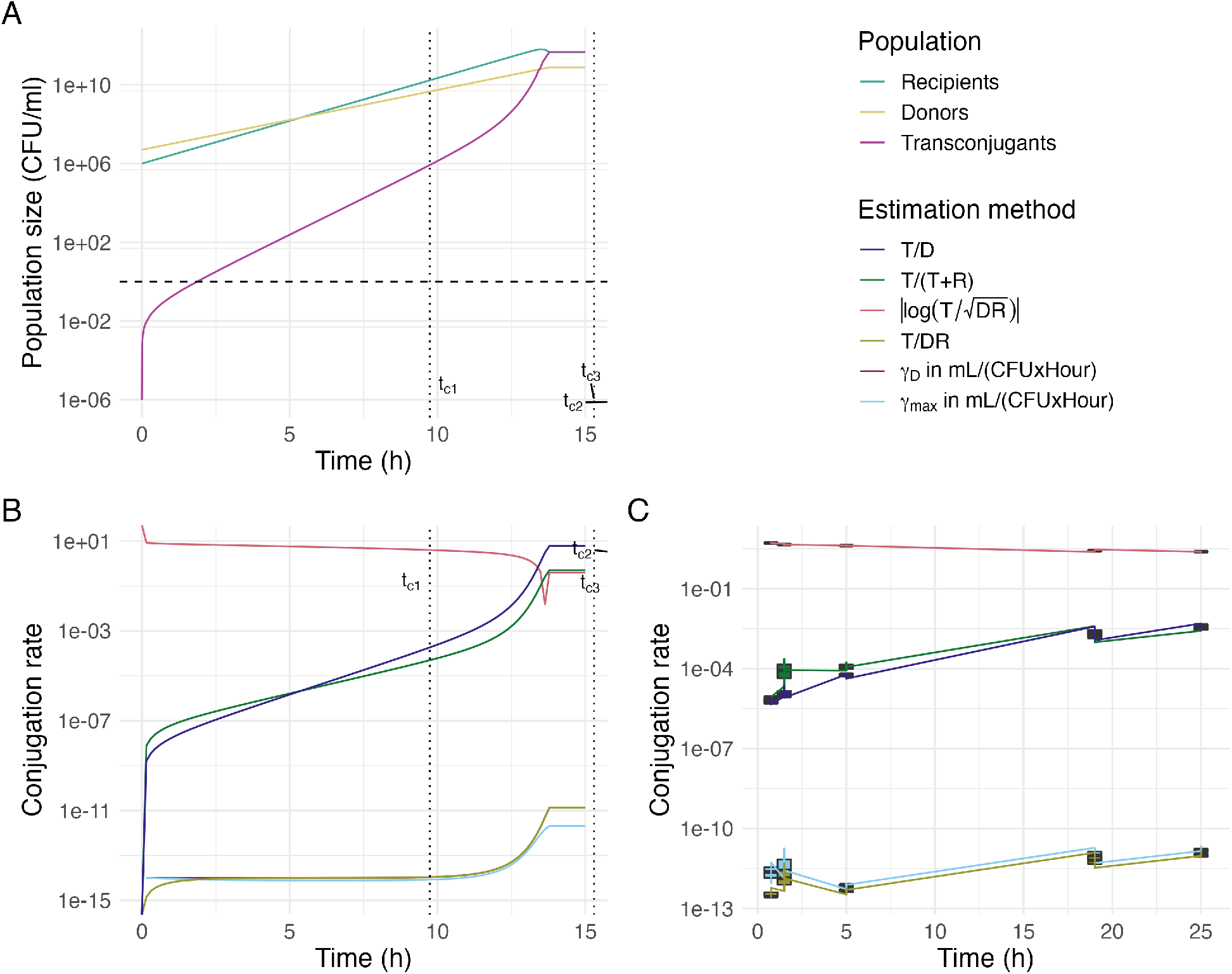
Impact of the time point of measurement on the magnitude of conjugation rate estimates. Panel **A** shows the simulated population dynamics; panel **B** shows the corresponding conjugation proficiency according to several population density and population dynamics based methods; panel **C** shows some of the same methods applied to data from a time course conjugation experiment. In panel **A,B** the simulation parameters were chosen to illustrate a cost of plasmid carriage, and a higher rate of conjugation from transconjugants to recipients than from donors. The SM estimate is denoted by *γ_max_* and the ASM estimate by *γ_D_*. The T/DR and *γ_D_* methods are partially overlaid. The vertical dotted lines indicate the first critical time, *t*_*c*1_ (at which the contribution of conjugation events from transconjugants becomes substantial), and the third critical time *t*_*c*3_ (see Supplementary Materials). In panel **A**, the horizontal dashed line indicates a single cell. The simulation parameters were: initial population densities *R*_0_ = 1 · 10^6^ CFU/mL, *D*_0_ = 5 · 10^6^ CFU/mL; initial resource concentration *C*_0_ = 10^12^ *μ*g/mL; growth rates *ψ_T_* = *ψ_D_* = 0.7, *ψ_R_* = 1.0 h^−1^; conjugation rates *γ_D_* = 10^−14^ mL · CFU^−1^h^−1^, *γ_T_* = 10^−11^ mL · CFU^−1^h^−1^; approximation factor *f* = 10. In panel **C**, we incubated a 1:1 mixture of donor D with recipient R in LB medium (n = 3), starting from an initial density of 1.5 · 10^7^ CFU/mL. Note that n = 2 at t = 19, due to an error in selective antibiotic plating. The measured growth rates were *ψ_D_* = 1.46, *ψ_R_* = 1.43, and ψ_T_ = 1.40.

As an example of the population-density based measures, we investigate the behaviour of the *T/D* method on a wider range of simulated data. Figure 2 shows that *T/D* varies multiple orders of magnitude as a function of the initial population densities and donor to recipient ratios. This variation occurs regardless of the measurement time point. If the initial population densities are manipulated, but the ratio of D:R is kept constant at 1:1 (Figure 2A), the *T/D* measure increases roughly proportional to the increased initial population density. Instead, if the total population density is kept constant, but the relative ratios of recipient and donor densities are varied (Figure 2B), the *T/D* measure declines roughly proportional to the change in initial recipient population density. The sensitive dependence on initial population densities, donor to recipient ratios, and time of measurement complicate the interpretation of population density based measures such as *T/D*. It also means that experimental condition that affect the initial donor and recipient population densities or ratios, will affect *T/D* independent of any true effects of the experimental condition on the plasmid conjugation rate.

**Figure 2:**
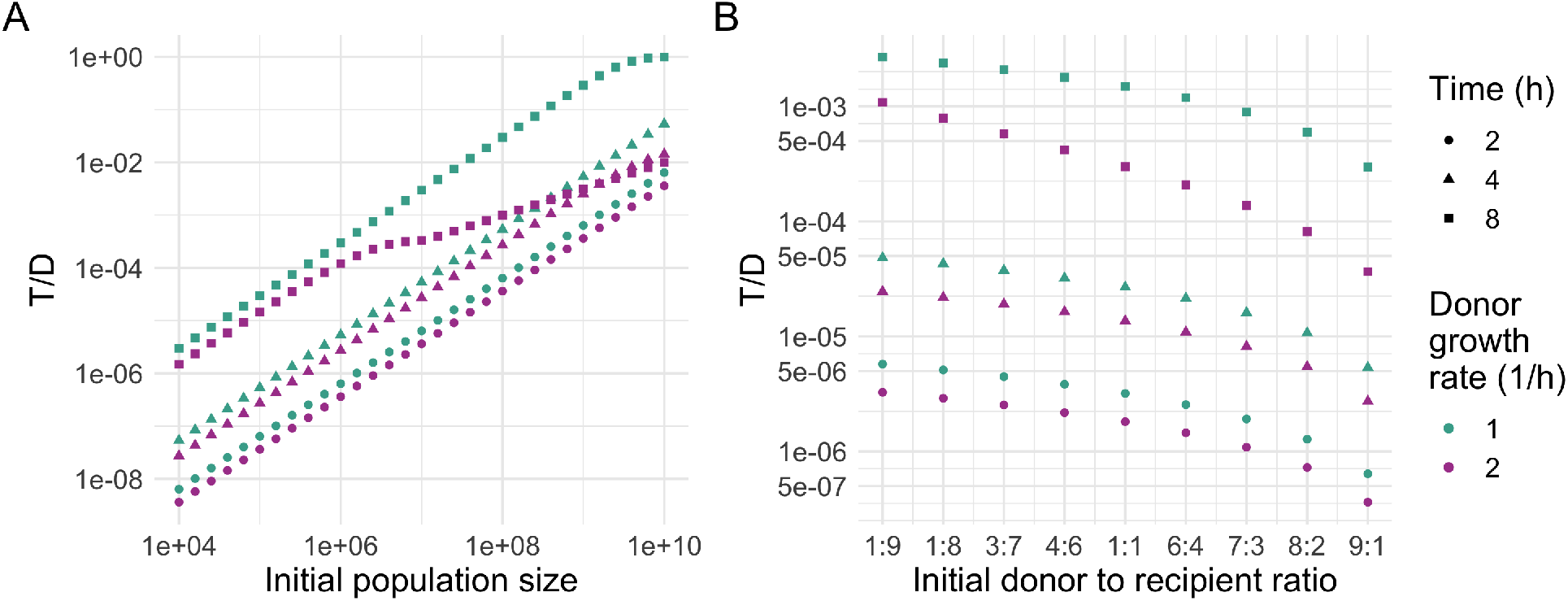
Impact of initial population density **(A)**, and donor-recipient ratio **(B)** on the *T/D* conjugation frequency estimate. The estimate varies over several orders of magnitude as a function of the initial population densities, relative population densities, and measurement time point. Parameters: initial resource *C*_0_ = 10^14^ *μ*g/mL; growth rates *ψ_T_* = *ψ_D_* = *ψ_R_* = 1.0 h^−1^; conjugation rates *γ_T/D_* = 10^−13^ mL · CFU^−1^h^−1^. For panel **(A)**, the initial population densities are *D*_0_, *R*_0_ ∈ [10^4^, 10^8^] CFU/mL. Recipient and donor populations are kept at the same density. For panel **(B)**, the ratio between initial population densities is *D*_0_: *R*_0_ ∈ [9:1, 1:9], with the total population density constant at 10^7^ CFU/mL.

Similarly, an experimental treatment that affects growth rates may confound the conclusion of the effect of treatment on conjugation rates, depending on the method used to estimate these rates (Fig. 3). A researcher may for instance wish to study the effect of sublethal concentrations of antibiotics, temperature, or nutrient conditions on plasmid conjugation rates, or establish whether a plasmid transfers with a higher rate to strain A than strain B. In each of these cases, the growth rate of donors, recipients, or transconjugants may be different in the treatment than the control. Especially population density based methods will conflate these growth rate differences with conjugation and may find significant effects of treatment on plasmid conjugation rates in the absence of any real effect (Fig. 3).

**Figure 3:**
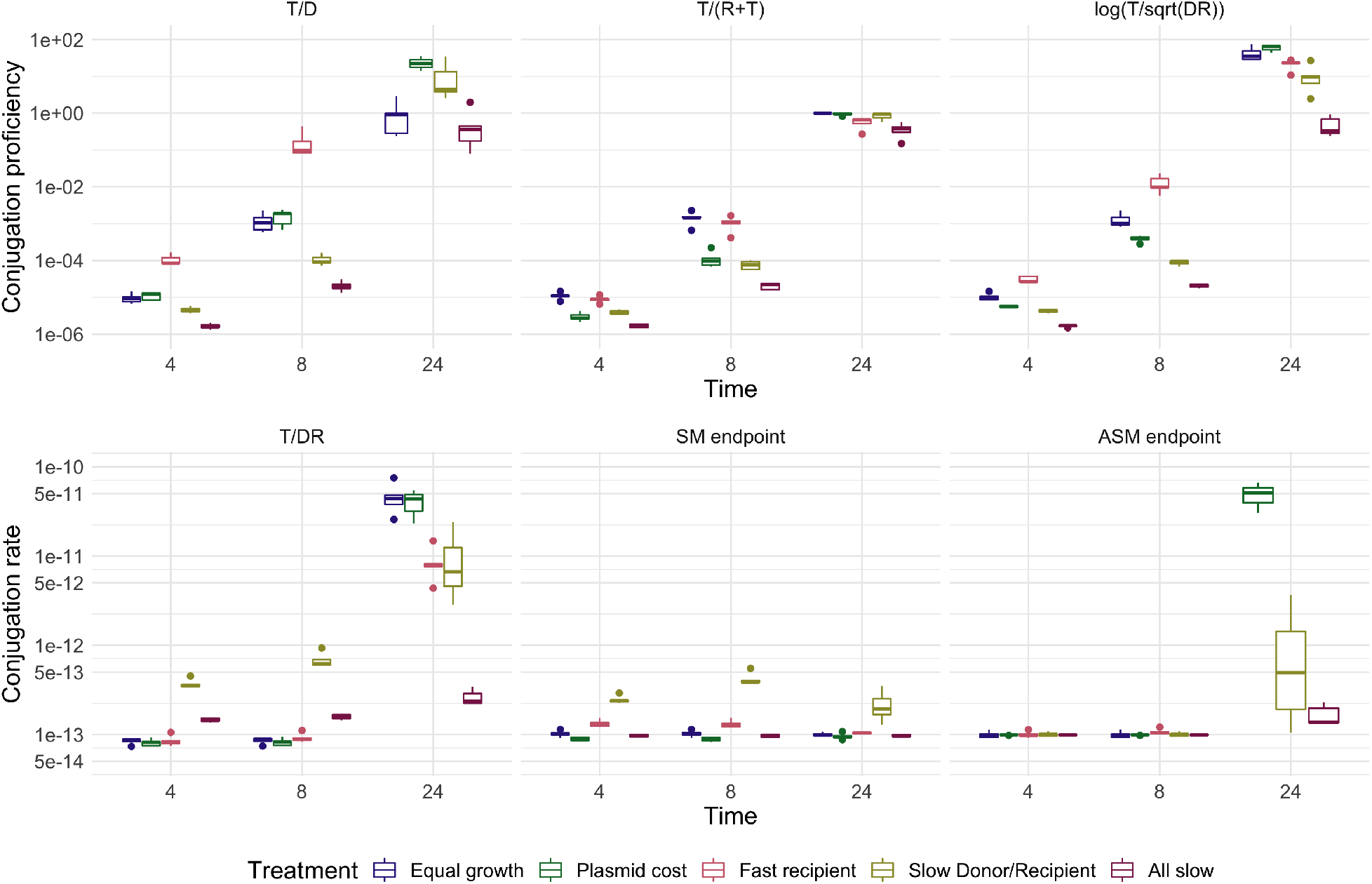
The effect of experimental treatments that affect growth rates, on conjugation rates estimated from simulated data. The true plasmid conjugation rate is drawn from a normal distribution to simulate observation errors, but otherwise kept the same across all panel and treatments 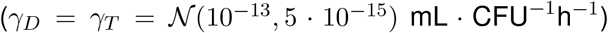. We compare a control (i) and 4 experimental scenarios (ii-v). (i) Equal growth: all strains grow equally fast at *ψ* = 1.2 h^−1^. (ii) Plasmid cost: donors and transconjugants have a 25% growth cost due to plasmid carriage. (iii) Fast recipient: the recipients and transconjugants grow 50% faster, e.g. because they are a different species. (iv) Slow Donor/Recipient: the growth rate of donor and recipient strains is reduced by 50%, e.g. due to antibiotic pre-treatment. (v) All slow: all populations grow 50% slower than in the control scenario (i), e.g. due to a different temperature or growth medium. The critical time of treatment (i) is 11-12 hours. We added 5% normal distributed noise to the model parameters to simulate experimental variation. For all treatments, the initial resource concentration *C*_0_ = 10^14^ *μ*g/mL, and initial densities 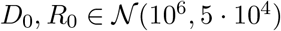 CFU/mL.

### Extending the Simonsen method

We have seen that population-based measures are not robust to variation in (i) the time-duration of the assay (Figure 1B), (ii) initial population densities (Figure 2A), (iii) and donor to recipient ratios (Figure 2B). The ‘end-point’ method based on the Simonsen model (SM), which has been around for 30 years, is robust to these factors. However, this method is not applicable to populations with differing growth rates (Fig. 3), nor differences in conjugation rates from donors and transconjugants. Thus, we extended the SM for differing growth and conjugation rates (see Materials and Methods and Supplementary Materials), and derived a similar ‘end-point’ formula for this new model (the ASM), which is easily computed on experimental data.

In deriving the ASM estimate, we make some assumptions about the relative size of different processes contributing to the overall dynamics of *D, R* and *T* populations. Some of these assumptions are also tacitly made in the SM estimate. Most prominently, this includes the assumptions that (i) the recipient population is not substantially reduced due to transformation to transconjugants, and (ii) no conjugation takes place in stationary phase. If the rates of conjugation from donors and transconjugants differ, both the SM and ASM further require that (iii) the populations were measured at a time where the dynamics are still dominated by conjugation events between donors and recipients rather than between transconjugants and recipients. When these assumptions are no longer valid, we expect the SM and ASM estimates for the donor conjugation rate to fail. By making these assumptions explicit, we can derive the critical time *t_crit_* beyond which the approximations break down (see the Supplementary Materials). Importantly, this critical time *t_crit_* is the minimum of three different time points, reached when one of the approximations (i) or (iii) fails. Which of these time points is reached first, and thus which dictates the latest possible measurement time point, depends on the relative magnitude of the growth rates (*ψ_D_, ψ_R_, ψ_T_*), conjugation rates (*γ_D_, *γ_T_**), as well as the initial population densities (*D*_0_, *R*_0_, see Supplementary Materials eqs. 9.24, 9.25, 9.26). There is some circularity in the expressions, since knowledge of the conjugation rates is required to determine when to measure the conjugation rates. This should become part of the routine of testing and setting up a new conjugation assay, and will not change much for strains with similar growth and conjugation rates. With species like *E. coli*, we generally recommend to measure early (e.g. after 4-8 hours) rather than to wait for overnight cultures.

We use simulated data to investigate whether the ASM estimate improves the conjugation rate estimate in the face of differing (i) growth, and (ii) conjugation rates. Here, we use the fold change, i.e. the ratio between the estimated value and the true value of the conjugation rate *γ_max_*, to quantify the error made during estimation.

### Growth

As Figure 4 shows, the SM estimate varies as a function of the donor and recipient population growth rate. The SM overestimates the conjugation rate if donor and/or recipients populations grow more slowly than the transconjugant population (lower left corner of Figure 4C). If the transconjugants grow more slowly than D and/or R, the SM underestimates the conjugation rate (upper right corner of Figure 4C). This is the case for all measurement time points, although the effect is exacerbated for measurements that are made after a longer conjugation time (Figure 4A). In contrast, the new ASM estimate *γ_D_* is valid until the critical time *t_crit_*, i.e. the time point for which the approximations of the model break down (Figure 4E). The critical time window grows shorter as the absolute magnitude of the growth rates increases (Figures 4B, D, and S1). Because the critical time is determined as the minimum of three different processes, all of which depend on the growth rates in different ways, the process dictating the critical time changes as a function of the growth rate. In Figure 4B the limiting process for low donor growth rates is the early onset of substantial conjugation from transconjugants (time *t*_*c*1_, see Supplementary Materials) and at higher donor growth rates the substantial reduction of the recipient population due to conjugation events (time *t*_*c*2_, see Supplementary Materials).

**Figure 4:**
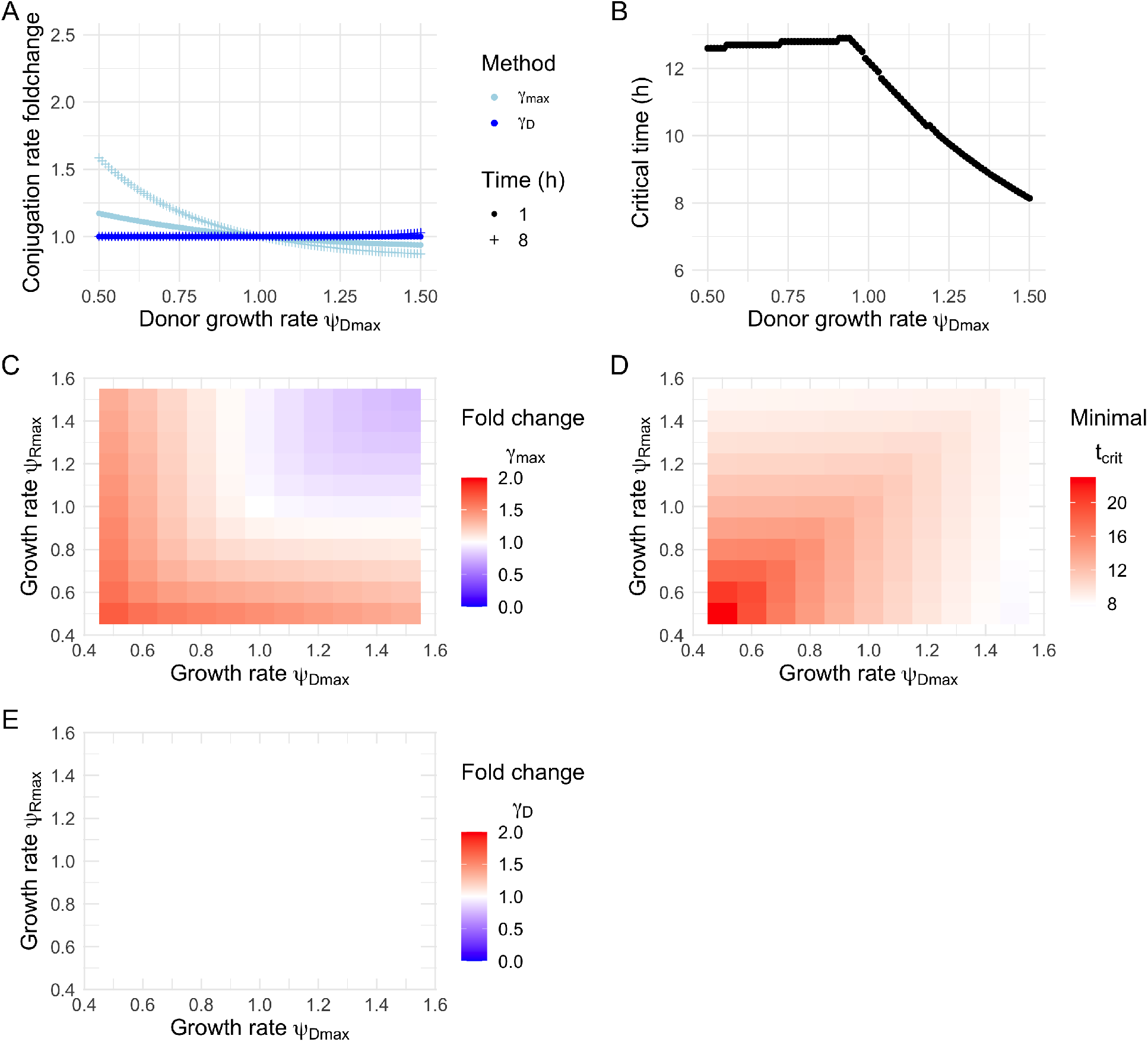
The effect of growth rate differences on the accuracy of the SM and ASM conjugation rate estimates. Panel **A** shows the deviation of the estimated conjugation rate from the true value (fold change 1) in the simulation as a function of the donor growth rate, where *γ_max_* denotes the SM estimate and *γ_D_* the ASM estimate (symbols denote different measurement time points). Panel **B** shows the corresponding critical time. Panels **C** and **E** show the same comparison to the true conjugation rate as panel **A**, but with additionally varying recipient growth rates and assuming measurement after 8 hours. Panel **E** seems empty because the ASM estimate is so close to the true value. Panel **D** shows the critical time corresponding to panels **C** and **E**. For deviating growth rates, the SM always shows a minor estimation error. Faster donor or recipient growth reduces the critical time, which is mirrored by the greater deviation of the estimated conjugation rates from the true value (especially for later measurement time points, as in **A**). Fold change is defined as the ratio between the estimated value and the true value. Parameters: initial population densities *R*_0_ = *D*_0_ = 5 · 10^6^ CFU/mL; initial resources *C*_0_ = 10^14^ *μ*g/mL; growth rate *ψ_T_* = 1.0 h^−1^; conjugation rates *γ_D_* = *γ_T_* = 10^−13^ mL · CFU^−1^h^−1^; approximation factor *f* = 10 are the same for all panels. For panels **A, B**, growth rate *ψ_D_* ∈ [0.5, 1.5] h^−1^, *ψ_R_* = 1 h^−1^. For panels **C, D, E**, growth rates *ψ_D_, ψ_R_* ∈ [0.5, 1.5] *h*^−1^.

### Conjugation rates

If the rates of conjugation from donors and transconjugants differ, both the SM and the ASM estimates accurately estimate the donor to recipient conjugation rate, as long as D, R, T are measured sufficiently early (Figure 5A/C/E). This is because the contribution of TRT conjugation events will be small as long as the transconjugant population is still small. For later times, the estimated SM conjugation rate *γ_max_* will interpolate between *γ_D_* and *γ_T_*. The estimated time at which the approximations break down (*t_crit_*) is the same for both methods (Figure 5B/D). As can be seen in Figures 5A and 5C/E, this means that the magnitude of the misestimation of SM and ASM estimates depends strongly on the measurement time point. This shows that it is critically important not to measure too late.

**Figure 5:**
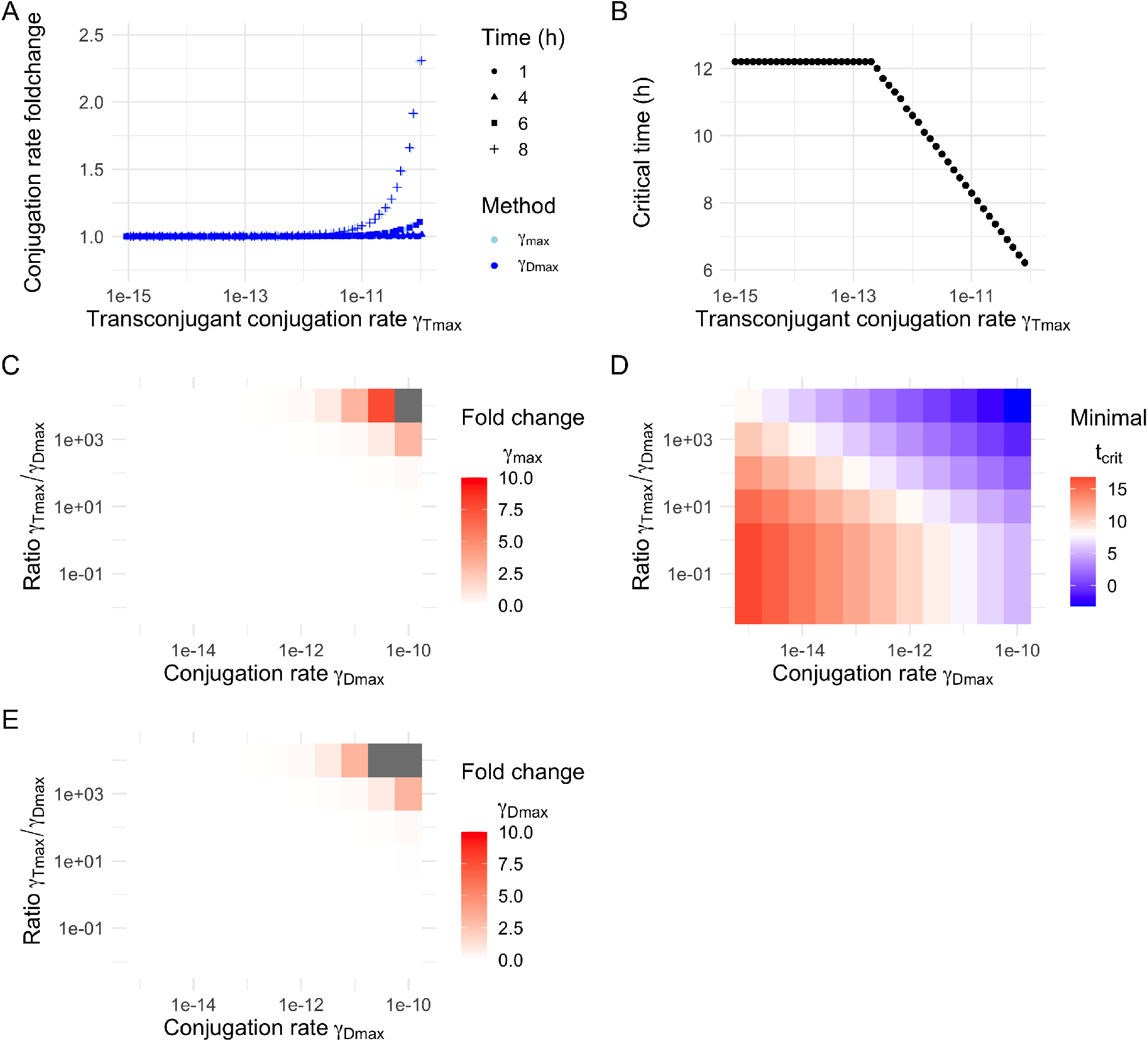
The effect of conjugation rate differences on the accuracy of the SM and ASM conjugation rate estimates. Panel **A** shows the deviation of the estimated conjugation rate from the true value (fold change 1) in the simulation as a function of the transconjugant conjugation rate, where *γ_max_* denotes the SM estimate and *γ_D_* the ASM estimate (symbols denote different measurement time points). Panel **B** shows the corresponding critical time. Panels **C, E** show the same comparison to the true conjugation rate as panel **A**, but with additionally varying donor conjugation rates and assuming measurement after 8 hours. Panel **D** shows the corresponding critical time. The ratio *γ_T_/γ_D_* **C, D, E** indicates how much bigger the rate of conjugation from transconjugants is than that from donors. For deviating transconjugant conjugation rates both methods are correct within the critical time **A** (the methods are partially overlaid). Faster conjugation rates reduce the critical time, which is mirrored by the greater deviation of the estimated conjugation rates from the true value (especially for later measurement time points, as in **A**). Fold change is defined as the ratio between the estimated value and the true value. Both the SM and ASM result in numerical errors when measuring substantially above the critical time, upper right corner of panels **C, E**. Parameters: initial population densities *R*_0_ = *D*_0_ = 5 · 10^6^ CFU/mL; initial resources *C*_0_ = 10^14^ *μ*g/mL; growth rates *ψ_D_* = *ψ_T_* = *ψ_R_* = 1.0 h^−1^; approximation factor *f* = 10 are the same for all panels. For panels **A, B**, conjugation rate *γ_D_* = 10^−13^ mL · CFU^−1^h^−1^, and conjugation rate *γ_T_* ∈ [10^−15^, 10^−10^] mL · CFU^−1^h^−1^. For panels **C, D, E**, conjugation rates *γ_D_* ∈ [10^−15^, 10^−10^] mL · CFU^−1^h^−1^, *γ_T_* ∈ [10^−17^, 10^−6^] mL · CFU^−1^h^−1^.

### Protocol

These theoretical considerations have led us to propose the following protocol to perform conjugation assays. In its most complete form the protocol requires two conjugation experiments: a first one starting from a *D* + *R* mixed culture, and then a second one with *T* + *R*′. The dash is to indicate that the recipients of the second experiment (*R*′; or the transconjugants from the first experiment) need to be provided with an additional selective marker such that the transconjugants of the second experiment (*T*′) can be distinguished from those of the first (*T*).

As pointed out in the previous section, it is important that the population densities of *D, R* and *T* are measured before the critical time is reached. Strictly speaking, this critical time can only be determined after both conjugation experiments are completed, as they require an estimate of both conjugation rates (*γ_D_, *γ_T_**), as well as all growth rates (*ψ_D_, ψ_R_, ψ_T_*, see Supplementary Materials eqs. 9.24, 9.25, 9.26). To optimise the chance of measuring below the critical time, we recommend to measure as soon as a measurable number of transconjugants has been formed.

Note, if one can assume that the difference between *γ_D_* and *γ_T_* is negligible, then the second conjugation experiment with *T* + *R*′ is not necessary. This protocol also does not capture the effect of transitory derepression and thus implicitly assumes that this state has a small effect on the conjugation rate from transconjugants compared to genetic factors. The general effect of transitory derepression would be to increase *γ_T_*, and thus reduce the critical time further. The R package and Shiny app we developed contains a function to determine how sensitively the minimal critical time depends on the presumed values of the conjugation rate from transconjugants *γ_T_*.

### Run 1st experiment with *D* and *R*

- Grow overnight cultures of *D* and *R*. We recommend diluting overnight cultures appropriately and growing strain cultures into early exponential phase before the start of the assay [4].
- The ASM method requires the initial densities of *D* and *R*, but is not sensitive to the exact value as long as the order of magnitude is correct (necessary to determine the critical time). Plating at *t*_0_ is therefore mostly optional.
- Incubate cultures of *D* and *R* in isolation and as a mixed culture of *D* + *R*. Measure the growth rates of all cultures in exponential phase. It can be convenient to set this experiment up in a plate reader and measure optical density through time. This yields estimates for the growth rates *ψ_D_* and *ψ_R_* (in h^−1^) from the single cultures, as well as *ψ_max_* from the mixed culture.
- Plate the mixed culture on selective plates at a time *t*_1_, to estimate the population densities of *D, R* and *T* (in CFU/mL). This time point should be early enough, such that there is a high chance that it is below the critical time *t*_*crit*,1_ for the 1st experiment. We recommend the inclusion of appropriate controls to test whether conjugation occurs on the double selective agar plate, rather than in liquid culture [9, 36, 37].
- Calculate the ASM estimate for the conjugation rate from donors *γ_D_*. This requires the initial population densities *D*_0_, *R*_0_; the growth rates *ψ_D_, ψ_R_, ψ_T_*; the time of measurement *t*_1_; and the measured population densities *D*_*t*1_, *R*_*t*1_, *T*_*t*1_.
- In case you are considering not to perform the 2nd conjugation experiment, you can use the R package or Shiny app to determine how sensitively the minimal critical time depends on the presumed values of the conjugation rate from transconjugants *γ_T_*.

### Run 2nd experiment with *T* and *R*′

- Isolate single transconjugant clones *T* from the 1st experiment, to use as plasmid donors in the 2nd experiment. Either these clones or the recipients used in this 2nd experiment need to be provided with an additional selective marker such that the transconjugants of the 2nd experiment (*T*′) can be distinguished from those of the 1st experiment (*T*).
- Grow overnight cultures of *T* and *R*′. We recommend diluting overnight cultures appropriately and growing strain cultures into early exponential phase before the start of the assay [4].
- Incubate cultures of *T* and *R*′ in isolation and as a mixed culture of *T* + *R*′. Measure the growth rates of all cultures in exponential phase. This yields estimates for the growth rates *ψ_T_* from the single cultures, as well as *ψ_max_* from the mixed culture.
- Plate the mixed culture on selective plates at a time *t*_2_, to estimate the population densities of *T, R*′ and *T*′. This time point should be early enough, such that there is a high chance that it is below the critical time *t*_*crit*,2_ for the 2nd experiment.
- Estimate the conjugation rate from transconjugants *γ_T_*.
- Check whether *t*_2_ < *t*_*crit*,2_ for the 2nd experiment.
- If the 2nd experiment is within the critical time, check whether *t*_1_ < *t*_*crit*,1_ for the 1st experiment.

If either *t*_1_ or *t*_2_ are too large, the experiments will need to be repeated, choosing times smaller than *t_crit_*.

In Figure 6 we show the results of such a full protocol for a conjugation experiment between two *E. coli* strains with similar growth rates. The conjugation rate from transconjugants was about one order of magnitude higher than from donors. The ASM estimate could not be computed based on the measurements taken after 24 hours, since they were past the critical time (9.3 hours). The SM estimate shows slightly higher *γ_T_* estimates (from the TRT experiment) after 24 hours than after 4, but this difference is not significant. The reason these estimates do not differ more strongly is likely because the stationary phase was reached relatively early (after 6 hours).

**Figure 6:**
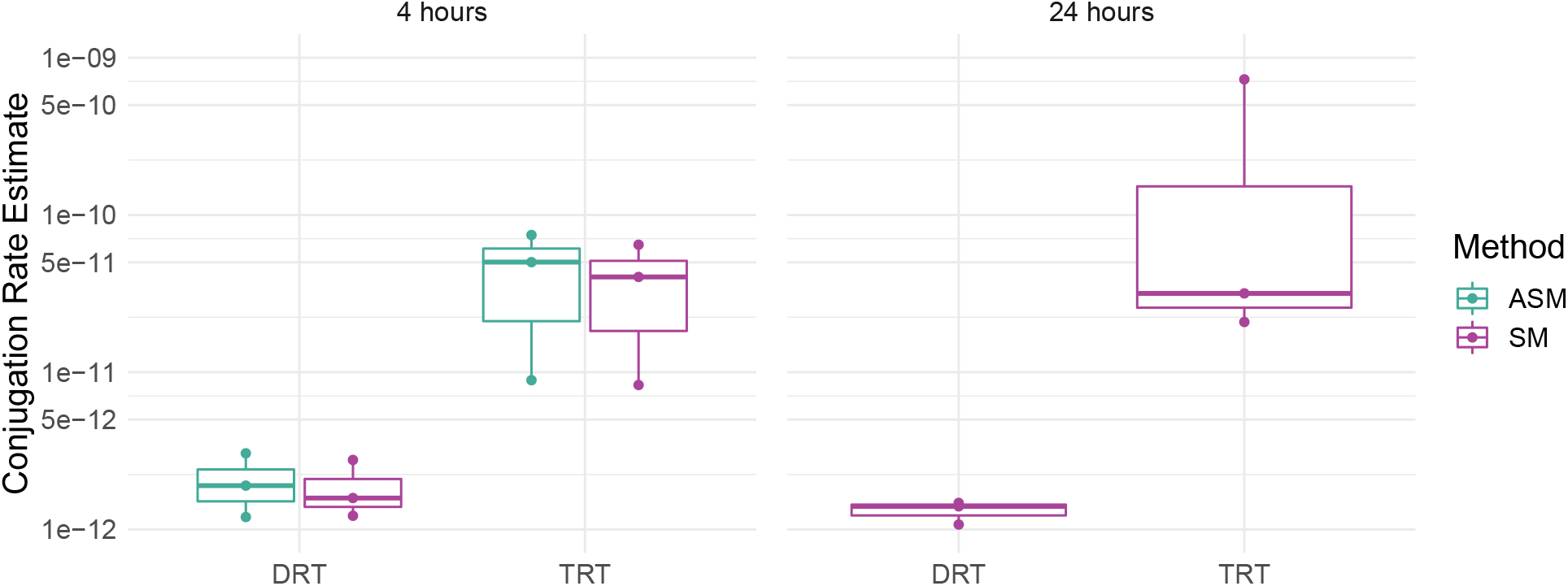
Full protocol measured after 4 and 24 hours, and conjugation rates estimated with the SM and ASM. The conjugation rate from transconjugants was about an order of magnitude higher than from donors. For these strains the minimal critical time was 9.3 hours, but stationary phase was reached already after 6. Donor D was mated with recipient R in VL medium (n = 3), according to the full protocol (see Methods). For the TRT experiment, we used a previously isolated transconjugant strain with spontaneous nalidixic acid resistance (D’). The Donors and recipients were mixed in a 1:1 ratio, starting at an initial density of 1 · 10^6^ CFU/mL.

## 5 Tools for the scientific community

We present an R package (https://github.com/JSHuisman/conjugator) which allows researchers to calculate various plasmid conjugation rates from experimental data, and check whether a given experiment was measured within the critical time. Currently the package includes the SM, ASM, *T/D, T/DR, T*/(*R* + *T*), 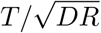, and 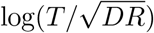. The package can be extended by other methods in the future, including recently proposed fluctuation test based methods [26].

To further enhance the accessibility of these functions, we added a graphical user interface to the package in the form of a Shiny web application (https://ibz-shiny.ethz.ch/jhuisman/conjugator/). This app allows researchers to (i) upload their own data, use the functions from the package, and download the results, as well as (ii) simulate bacterial population dynamics with conjugation to get a better feeling for how the different conjugation measures behave depending on different parameters of the experiment.

## 6 Discussion

There is no gold standard to determine and report conjugation rates, and this has complicated the comparison of experimental values obtained by different research groups or under different (a)biotic conditions [15, 40]. We have presented an overview of the different methods found in the literature, and exemplified how commonly used methods are affected by initial population densities, donor to recipient ratios, differences in growth rate of the mating strains, or measurement time point. As much as possible, we propose to settle on a single method to describe conjugation proficiency [19]. Ideally, such a measure would allow comparison across experimental conditions, and to parametrise mechanistic models used to explain and predict plasmid dynamics. Both the SM and ASM methods are reasonable for this purpose.

If the donor, recipient, or transconjugant populations differ in their growth rates, the SM makes a minor estimation error that is corrected by using our new ASM estimate. When the conjugation rate from transconjugants differs substantially from the donor conjugation rate, both methods estimate a correct conjugation rate only during the initial phase of the experiment, before the critical time is reached. Overall, we find that bacteria with large growth rate differences, high absolute growth rates, and high absolute conjugation rates are most likely to lead to problems in conjugation rate estimation, as these factors speed up the population dynamics and reduce the critical time.

To encourage ‘best practices’ in the estimation and reporting of conjugation rates, we developed an R package and web application that compute these values from experimental data. A further clear conclusion of this work is that one should measure the outcome of conjugation assays early, before the dynamics become dominated by conjugation from transconjugants. Our critical time estimates give an indication of how early this should be.

Several caveats remain for both the SM and the ASM. First, these models are in principle not suitable for application to mating assays on solid surfaces, as they assume well-mixed conjugating populations. However, the conjugation rates in high-density, well-mixed surface mating experiments are comparable to liquid mating, provided they are measured sufficiently early [18]. Second, the ASM assumes that the growth rates in monoculture are predictive for the same strains in mating populations, and thus disregards competitive effects. Direct measurements of the individual mixed culture growth rates would require *in situ* tracking of donors, recipients and transconjugants via differential fluorescent labelling of the plasmid and the recipient chromosome. Last, these methods are based on population dynamic models that assume no dependence of conjugation on cell density, no induction in stationary phase, and no segregational loss. These assumptions may hold for IncF plasmids, but do not extend to all plasmid families [41, 42]. Future work should establish how common such ‘atypical’ plasmids are, and develop methods to quantify the conjugation rate also in these cases. Until then, these concerns could be addressed by constructing a more complex conjugation and growth model and fitting it to time course data of the different mating populations [20, 27, 35]. Further methods can be added to the conjugator R package in the future.

## 7 Acknowledgements

We would like to thank the members of the Theoretical Biology and Pathogen Ecology groups for helpful discussions, and Philip Ruelens (Wageningen University) and Mark Zwart (The Netherlands Institute of Ecology (NIOO-KNAW), Wageningen, The Netherlands) for scripts used in the experimental data analysis. This work was supported by the Swiss National Science Foundation (grant 407240-167121), and ZonMw (grant 541001005).

## Author Contributions

- JSH: Conceptualization, Formal analysis, Investigation, Methodology, Software, Validation, Visualization, Writing - original draft, Writing - review & editing
- FB: Conceptualization, Validation, Writing - review & editing
- SJND: Investigation, Validation, Writing - review & editing
- JAGMdV: Funding acquisition, Resources, Supervision, Writing - review & editing
- ARH: Conceptualization, Funding acquisition, Supervision, Writing - review & editing
- EAJF: Funding acquisition, Methodology, Software, Supervision, Writing - review & editing
- SB: Conceptualization, Funding acquisition, Methodology, Resources, Software, Supervision, Writing - review & editing

## 9 Supplementary Materials

### 9.1 ‘End-point’ method for the approximate extended Simonsen model

We aim to derive a simple formula to estimate the conjugation rate from donors, analogous to the SM ‘end-point’ method. To do so, we start from the equations for the approximate extended Simonsen model (ASM):

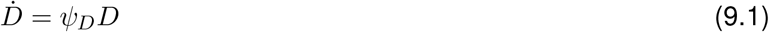

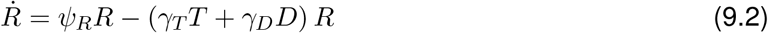

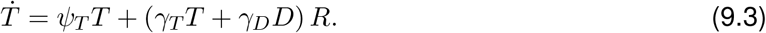

Assuming that (i) initially the recipient population dynamics are dominated by growth, and that (ii) the transconjugant population is not yet dominated by conjugation from transconjugants, i.e.:

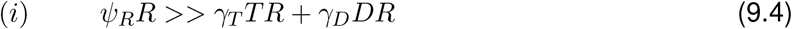

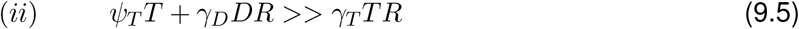

We obtain a simplified set of equations given by:

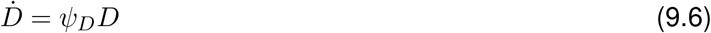

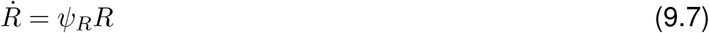

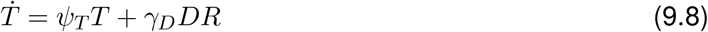

We solve this for initial conditions corresponding to an ‘invasion from rare’ scenario, i.e. *D*(0) = *D*_0_, *R*(0) = *R*_0_ and *T*(0) = 0, and get the solution:

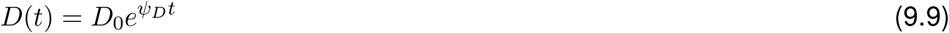

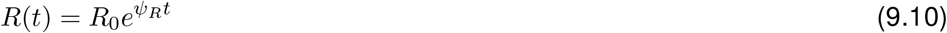

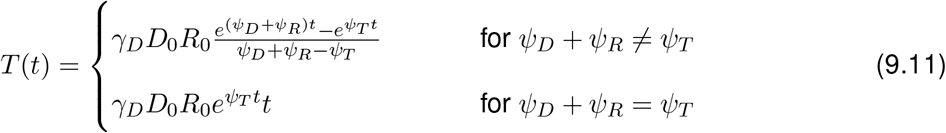

#### Conjugation rate

*γ_D_* This solution for *T* (eq. 9.11) contains the conjugation rate *γ_D_*. By rearranging the terms, and using equations 9.9 and 9.10 to substitute *D*(*t*)*R*(*t*) = *D*_0_*R*_0_*e*^(*ψ_D_*+*ψ_R_*)*t*^, we obtain an estimate of the conjugation rate *γ_D_* at time point *t*:

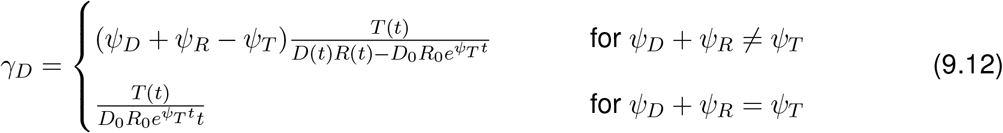

This expression (the ASM estimate) can be used instead of the SM estimate as long as the approximate equations (eqs. 9.6 - 9.8) are good approximations to the full ODE (eqs. 9.1 - 9.3). Since this solution requires that *D*(*t*) ≈ *D*_0_*e^ψ_D_t^* and *R*(*t*) ≈ *R*_0_*e^ψ_R_t^*, one really need only measure *T*(*t*) and the growth rates *ψ_D_, ψ_R_, ψ_T_*.

#### Critical time *t_crit_*

In deriving the ASM estimate, we made some approximations (eqs. 9.4 and 9.5) about the relative size of different processes contributing to the overall dynamics of *D, R* and *T* populations (leading to eqs. 9.9 - 9.11). When these approximations are no longer valid, the ASM estimate for *γ_D_* (eq. 9.12) fails. However, we can calculate the ‘critical time’ beyond which the approximations no longer hold.

First, the equation for 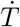 (eq. 9.8) fails to approximate the full ODE (eq. 9.3) once conjugation from transconjugants is substantial, i.e. once *γ_T_T*(*t*)*R*(*t*) ≈ *ψ_T_T*(*t*) + *γ_D_D*(*t*)*R*(*t*). If we specify a factor f by which the left hand side (conjugation from transconjugants) should be smaller than the right hand side (clonal growth of transconjugants and conjugation from donors), we obtain an equation for the time *t*_*c*1_ when the approximation will be violated:

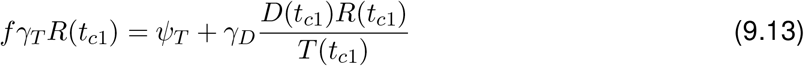

Here we already divided by *T*(*t*) on both sides. For the last term of this equation, we can substitute our approximation of *γ_D_* (eq. 9.12; for *ψ_D_* + *ψ_R_* ≠ *ψ_T_*) to obtain:

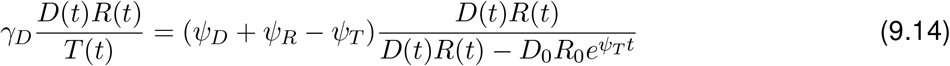

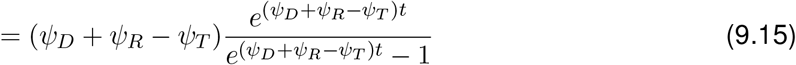

If we then assume that *t* >> 1/(*ψ_D_* + *ψ_R_* – *ψ_T_*), i.e. time *t* is substantially larger than the bacterial doubling time, we see that:

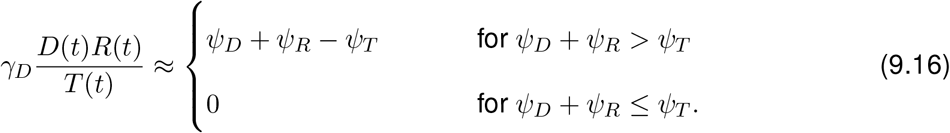

Substituting this expression (eq. 9.16) into equation 9.13 we get:

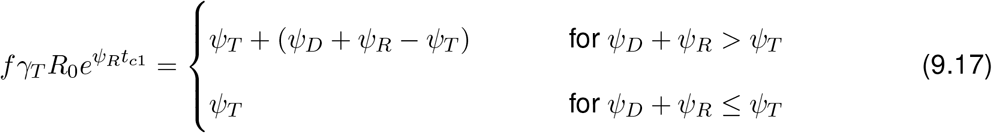

and thus, for the first critical time:

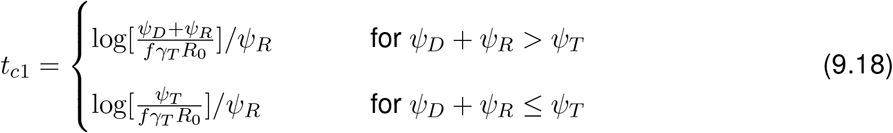

Second, 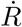 (eq. 9.7) fails to approximate the full ODE (eq. 9.2) once the recipient population dynamics are no longer dominated by growth, i.e. *ψ_R_R*(*t*) ≈ *γ_D_D*(*t*)*R*(*t*) + *γ_T_T*(*t*)*R*(*t*). To simplify this we break the approximation down into two parts: (i) *ψ_R_* ≈ *γ_D_D*(*t*) and (ii) *ψ_R_* ≈ *γ_T_T*(*t*).

Substituting the above expression for *D* (eqs. 9.9) into the first equation we find the second critical time:

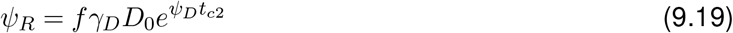

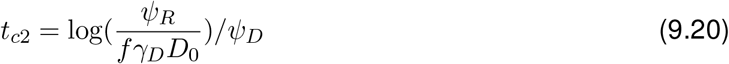

For the second equation we substitute the expression for *T*(*t*) (eq. 9.11) to get:

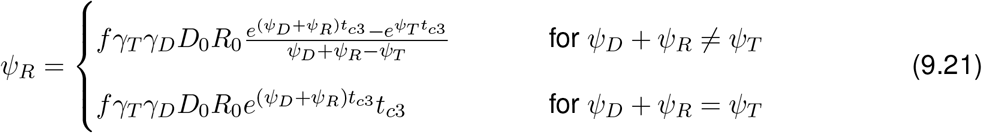

We again assume that the time *t* is substantially larger than the doubling time of the bacteria, i.e. *t* >> 1/(*ψ_D_* + *ψ_R_* – *ψ_T_*), to simplify it to:

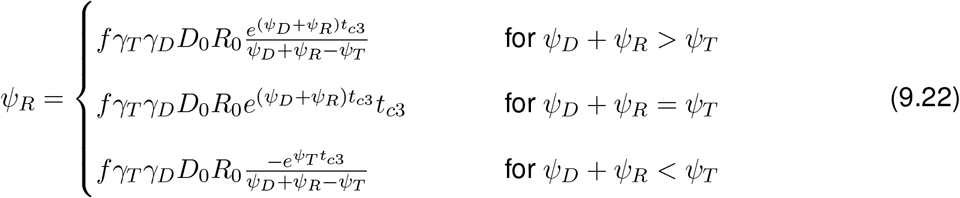

The equation 9.22 can not be solved analytically for the case where *ψ_D_* + *ψ_R_* = *ψ_T_*. In the R package this is solved numerically using a root finding algorithm. If we solve the other two cases of equation 9.22 for time, we obtain for the last critical time:

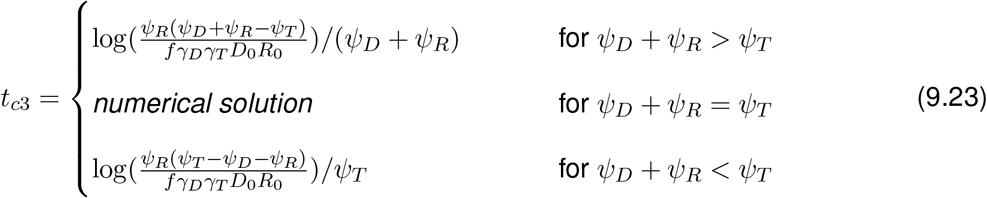

The ASM estimate will lose its validity as soon as one of the three critical times is reached. Depending on the parameters (the relative magnitude of growth and conjugation rates, as well as initial population densities) this could be any one of *t*_*c*1_ – *t*_*c*3_. With ‘the’ critical time *t_crit_*, we thus refer to the minimum *t_crit_* = min(*t*_*c*1_, *t*_*c*2_, *t*_*c*3_) of these three time points:

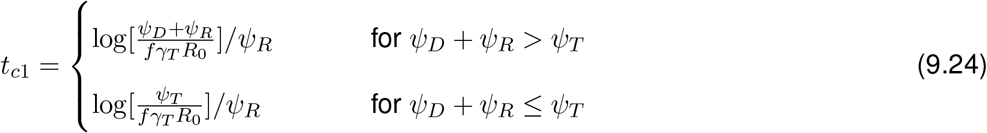

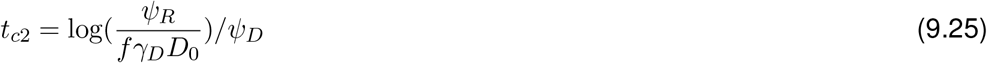

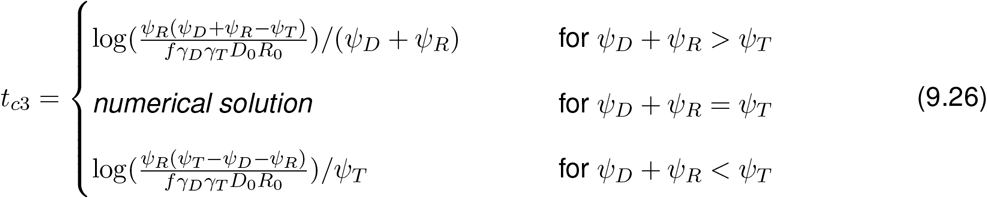

### 9.2 Stationary phase time

Stationary phase is reached at the time *t_stat_* when all initial resources *C*(*t* = 0) = *C*_0_ have been consumed by the growing bacteria, and converted into biomass; i.e.

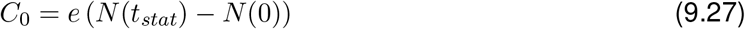

In a general case, *N*(*t*) = *D*(*t*) + *R*(*t*) + *T*(*t*) depends on the growth rate of all three populations, and the rate at which recipients are turned into transconjugants. If we substitute our earlier approximations for *D*(*t*), *R*(*t*) and *T*(*t*) we get:

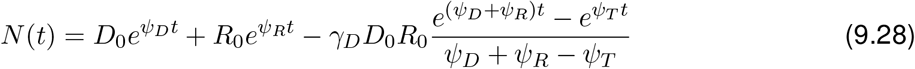

When all populations grow at the same rate *ψ_Xmax_*, any transformation of recipients into transconjugants does not affect the total population growth of N. If we assume simple exponential growth (as opposed to e.g. Monod dynamics), *N*(*t*) will be given by:

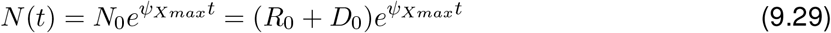

With this equation 9.27 becomes:

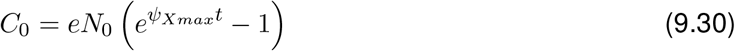

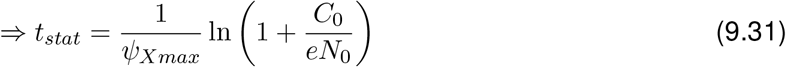

where *e* is in *μ*g per CFU. Population densities *N, D, T, R, N*_0_ in CFU/mL. Resource *C* in *μ*g per mL. Growth rate *ψ_Xmax_* is per hour.

In cases where we observe the mating population, we can simply replace *ψ_Xmax_* by the growth rate of that mixed population (*ψ_max_* from the Simonsen model). If one were to include Monod like growth dynamics, this would slow down growth at high population densities/ as the resource is becoming depleted. As a result, the start of stationary phase *t_stat_* would be slightly delayed.

### 9.3 The impact of higher conjugation rates on conjugation rate estimation

**Figure S1:**
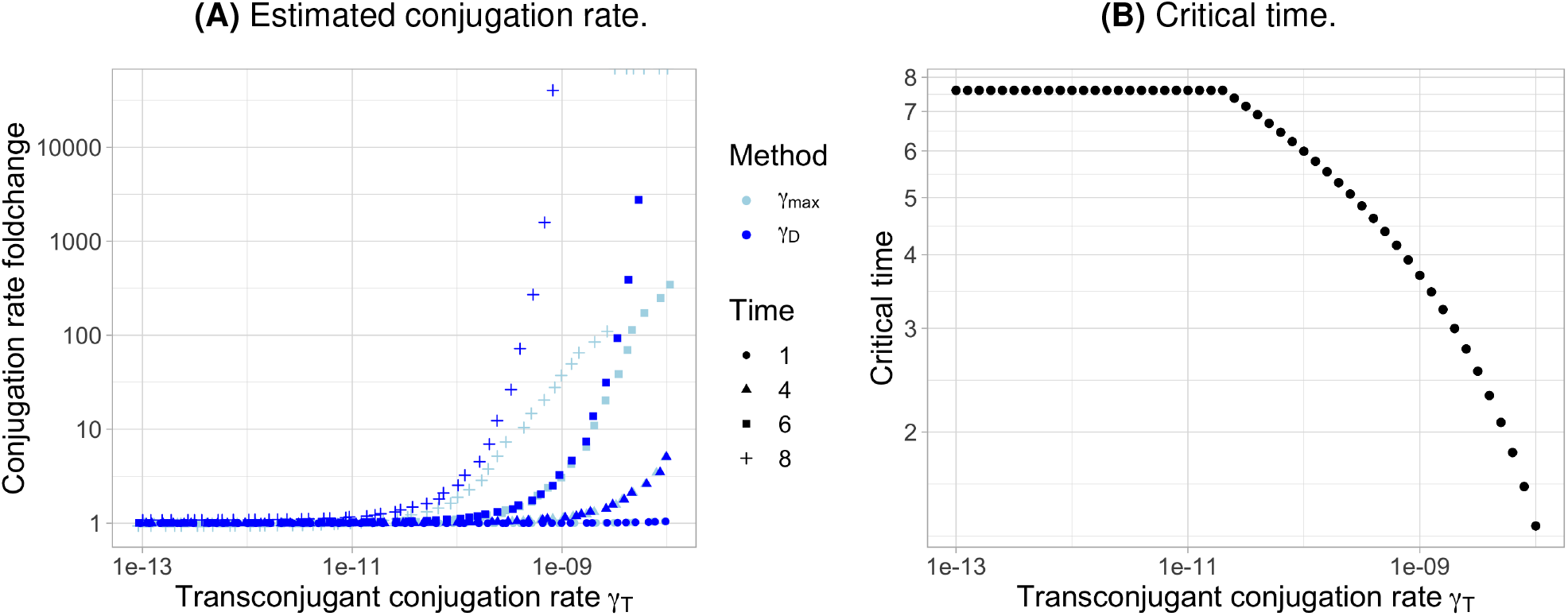
Panel **(A)** shows the ratio between the estimated conjugation rate and the true value in the simulation, for different measurement time points, as a function of the donor growth rate (*γ_max_* denotes the SM estimate, *γ_D_* the ASM estimate). Panel **(B)** shows the corresponding critical time. The parameters mimic those in Figure 5A/B in the main text, except that the conjugation rates are higher, and thus the temporal dynamics are faster. Initial population densities *R*_0_ = *D*_0_ = 5 · 10^6^ CFU/mL; initial resource concentration *C*_0_ = 10^14^ *μ*g/mL; growth rates *ψ_T_* = *ψ_D_* = *ψ_R_* = 1.0 h^−1^; conjugation rate *γ_D_* = 10^−11^ mL · CFU^−1^h^−1^; approximation factor *f* = 10.

## References

[1] Howard Ochman, Jeffrey G Lawrence, and Eduardo A Groisman. Lateral gene transfer and the nature of bacterial innovation. Nature, 405(6784):299, 2000. doi: 10.1038/35012500.

[2] James PJ Hall, Michael A Brockhurst, and Ellie Harrison. Sampling the mobile gene pool: innovation via horizontal gene transfer in bacteria. Philosophical Transactions of the Royal Society B: Biological Sciences, 372(1735):20160424, 2017. doi: 10.1098/rstb.2016.0424.

[3] Christian J.H. Von Wintersdorff, John Penders, Julius M. Van Niekerk, Nathan D. Mills, Snehali Majumder, Lieke B. Van Alphen, Paul H.M. Savelkoul, and Petra F.G. Wolffs. Dissemination of antimicrobial resistance in microbial ecosystems through horizontal gene transfer. Frontiers in Microbiology, 7(Feb):1–10, 2016. ISSN 1664302X. doi: 10.3389/fmicb.2016.00173.

[4] Allison J Lopatkin, Shuqiang Huang, Robert P Smith, Jaydeep K Srimani, Tatyana A. Sysoeva, Sharon Bewick, David K. Karig, and Lingchong You. Antibiotics as a selective driver for conjugation dynamics. Nature Microbiology, 1(6):16044, June 2016. ISSN 2058-5276. doi: 10.1038/nmicrobiol.2016.44.

[5] D. J. Rankin, E. P.C. Rocha, and S. P. Brown. What traits are carried on mobile genetic elements, and why. Heredity, 106(1):1–10, 2011. ISSN 0018067X. doi: 10.1038/hdy.2010.24.

[6] José Luis Martínez. Ecology and Evolution of Chromosomal Gene Transfer between Environmental Microorganisms and Pathogens. Microbiology Spectrum, 6(1):1–16, 2018. ISSN 2165-0497. doi: 10.1128/microbiolspec.mtbp-0006-2016.

[7] P. D. Lundquist and B. R. Levin. Transitory derepression and the maintenance of conjugative plasmids. Genetics, 113(3):483–497, 1986. ISSN 00166731. doi: 10.1093/genetics/113.3.483.

[8] Laura S. Frost and Günther Koraimann. Regulation of bacterial conjugation: balancing opportunity with adversity. Future Microbiology, 5(7):1057–1071, jul 2010. ISSN 1746-0913. doi: 10.2217/fmb.10.70.

[9] Fabienne Benz, Jana S Huisman, Erik Bakkeren, Joana A Herter, Tanja Stadler, Martin Ackermann, Médéric Diard, Adrian Egli, Alex R Hall, Wolf-Dietrich Hardt, and Sebastian Bonhoeffer. Plasmid-and strain-specific factors drive variation in ESBL-plasmid spread in vitro and in vivo. The ISME Journal, 15(3):862–878, 2021. ISSN 1751-7370. doi: 10.1038/s41396-020-00819-4.

[10] Tatiana Dimitriu, Lauren Marchant, Angus Buckling, and Ben Raymond. Bacteria from natural populations transfer plasmids mostly towards their kin. Proceedings of the Royal Society B: Biological Sciences, 286(1905):20191110, jun 2019. ISSN 14712954. doi: 10.1098/rspb.2019.1110.

[11] Amy J Mathers, Gisele Peirano, and Johann D.D. Pitout. The role of epidemic resistance plasmids and international high-risk clones in the spread of multidrug-resistant Enterobacteriaceae. Clinical Microbiology Reviews, 28(3):565–591, 2015. ISSN 10986618. doi: 10.1128/CMR.00116-14.

[12] Ricardo León-Sampedro, Javier DelaFuente, Cristina Díaz-Agero, Thomas Crellen, Patrick Musicha, Jerónimo Rodríguez-Beltrán, Carmen de la Vega, Marta Hernández-García, Nieves López-Fresneña, Patricia Ruiz-Garbajosa, Rafael Cantón, Ben S. Cooper, and Álvaro San Millán. Pervasive transmission of a carbapenem resistance plasmid in the gut microbiota of hospitalized patients. Nature Microbiology, 6(5):606–616, 2021. ISSN 20585276. doi: 10.1038/s41564-021-00879-y. URL http://dx.doi.org/10.1038/s41564-021-00879-y.

[13] Lone Simonsen, D. M. Gordon, F. M. Stewart, and Bruce R. Levin. Estimating the rate of plasmid transfer: an end-point method. Journal of General Microbiology, 136(11):2319–2325, 1990. ISSN 0022-1287. doi: 10.1099/00221287-136-11-2319.

[14] Cecilia Dahlberg, Maria Bergström, and Malte Hermansson. In situ detection of high levels of horizontal plasmid transfer in marine bacterial communities. Applied and Environmental Microbiology, 64(7):2670–2675, 1998. ISSN 00992240. doi: 10.1128/aem.64.7.2670-2675.1998.

[15] Richard J. Sheppard, Alice E. Beddis, and Timothy G. Barraclough. The role of hosts, plasmids and environment in determining plasmid transfer rates: A meta-analysis. Plasmid, 108(March), 2020. ISSN 10959890. doi: 10.1016/j.plasmid.2020.102489.

[16] Erik Bakkeren, Jana S Huisman, Stefan A Fattinger, Annika Hausmann, Markus Furter, Adrian Egli, Emma Slack, Mikael E Sellin, Sebastian Bonhoeffer, Roland R Regoes, Médéric Diard, and Wolf Dietrich Hardt. Salmonella persisters promote the spread of antibiotic resistance plasmids in the gut. Nature, 573(7773):276–280, sep 2019. ISSN 14764687. doi: 10.1038/s41586-019-1521-8.

[17] Fiona Flett, Vassilios Mersinias, and Colin P. Smith. High efficiency intergeneric conjugal transfer of plasmid DNA from Escherichia coli to methyl DNA-restricting streptomycetes. FEMS Microbiology Letters, 155(2):223–229, 1997. ISSN 03781097. doi: 10.1016/S0378-1097(97)00392-3.

[18] Xue Zhong, Jason Droesch, Randal Fox, Eva M. Top, and Stephen M. Krone. On the meaning and estimation of plasmid transfer rates for surface-associated and well-mixed bacterial populations. Journal of Theoretical Biology, 294:144–152, 2012. ISSN 00225193. doi: 10.1016/j.jtbi.2011.10.034.

[19] Søren J Sørensen, Mark Bailey, Lars H Hansen, Niels Kroer, and Stefan Wuertz. Studying plasmid horizontal transfer in situ: a critical review. Nature Reviews Microbiology, 3(9):700–710, 2005. doi: 10.1038/nrmicro1232.

[20] David Kneis, Teppo Hiltunen, and Stefanie Heß. A high-throughput approach to the culture-based estimation of plasmid transfer rates. Plasmid, 101:28–34, 2019. ISSN 10959890. doi: 10.1016/j.plasmid.2018.12.003.

[21] Marta Rozwandowicz, Michael S.M. Brouwer, Lapo Mughini-Gras, Jaap A. Wagenaar, Bruno Gonzalez-Zorn, Dik J. Mevius, and Joost Hordijk. Successful Host Adaptation of IncK2 Plasmids. Frontiers in Microbiology, 10(October):1–9, 2019. ISSN 1664302X. doi: 10.3389/fmicb.2019.02384.

[22] Gang Liu, Karolina Bogaj, Valeria Bortolaia, John Elmerdahl Olsen, and Line Elnif Thomsen. Antibiotic-Induced, Increased Conjugative Transfer Is Common to Diverse Naturally Occurring ESBL Plasmids in Escherichia coli. Frontiers in Microbiology, 10(September):1–12, 2019. ISSN 1664302X. doi: 10.3389/fmicb.2019.02119.

[23] Bruce R Levin, Frank M Stewart, and Virginia A Rice. The kinetics of conjugative plasmid transmission: fit of a simple mass action model. Plasmid, 2(2):247–260, 1979. doi: 10.1016/0147-619X(79)90043-X.

[24] Eva-Maria Saliu, Marita Eitinger, Jürgen Zentek, and Wilfried Vahjen. Nutrition Related Stress Factors Reduce the Transfer of Extended-Spectrum Beta-Lactamase Resistance Genes between an Escherichia coli Donor and a Salmonella Typhimurium Recipient In Vitro. Biomolecules, 9(8):324, jul 2019. ISSN 2218-273X. doi: 10.3390/biom9080324.

[25] P. Trieu-Cuot, C. Carlier, P. Martin, and P. Courvalin. Plasmid transfer by conjugation from Escherichia coli to Gram-positive bacteria. FEMS Microbiology Letters, 48, 1987. doi: 10.1111/j.1574-6968.1987.tb02558.x.

[26] Olivia Kosterlitz, Clint Elg, Ivana Bozic, Eva M. Top, and Benjamin Kerr. Estimating the rate of plasmid transfer with an adapted luria–delbrück fluctuation analysis and a case study on the evolution of plasmid transfer rate. bioRxiv, 2021. doi: 10.1101/2021.01.06.425583.

[27] Egil A.J. Fischer, Cindy M. Dierikx, Alieda Van Essen-Zandbergen, Herman J.W. Van Roermund, Dik J. Mevius, Arjan Stegeman, and Don Klinkenberg. The IncI1 plasmid carrying the bla CTX-M-1 gene persists in in vitro culture of a Escherichia coli strain from broilers. BMC Microbiology, 14(1):1–9, 2014. ISSN 14712180. doi: 10.1186/1471-2180-14-77.

[28] Roy Curtiss, Lucien G Caro, David P Allison, and Donald R Stallions. Early stages of conjugation in escherichia coli. Journal of bacteriology, 100(2):1091–1104, 1969. doi: 10.1128/jb.100.2.1091-1104.1969.

[29] Cagla Stevenson, James P J Hall, Michael A Brockhurst, and Ellie Harrison. Plasmid stability is enhanced by higher-frequency pulses of positive selection. Philosophical Transactions of the Royal Society B: Biological Sciences, 2018. doi: 10.1098/rspb.2017.2497.

[30] Catalina Arango Pinedo and Barth F Smets. Conjugal tol transfer from pseudomonas putida to pseudomonas aeruginosa: effects of restriction proficiency, toxicant exposure, cell density ratios, and conjugation detection method on observed transfer efficiencies. Appl. Environ. Microbiol., 71(1):51–57, 2005. doi: 10.1128/AEM.71.1.51-57.2005.

[31] Francisco Dionisio, Ivan Matic, Miroslav Radman, Olivia R Rodrigues, and François Taddei. Plasmids spread very fast in heterogeneous bacterial communities. Genetics, 162(4):1525–32, dec 2002. ISSN 1943-2631. doi: 10.1093/genetics/162.4.1525.

[32] João Alves Gama, Zilhão Rita, and Francisco Dionisio. Multiple plasmid interference - Pledging allegiance to my enemy’s enemy. Plasmid, 93(August):17–23, 2017. ISSN 10959890. doi: 10.1016/j.plasmid.2017.08.002.

[33] Sarah JN Duxbury, Jesse B Alderliesten, Mark P Zwart, Arjan Stegeman, Egil AJ Fischer, and J Arjan GM de Visser. Chicken gut microbiome members limit the spread of an antimicrobial resistance plasmid in *Escherichia coli*. Proceedings of the Royal Society B, 288(1962):20212027, 2021.

[34] Erik Gullberg, Lisa M. Albrecht, Christoffer Karlsson, Linus Sandegren, and Dan I. Andersson. Selection of a Multidrug Resistance Plasmid by Sublethal Levels of Antibiotics and Heavy Metals. mBio, 5(5):e01918–14, oct 2014. ISSN 2161-2129. doi: 10.1128/mBio.01918-14.

[35] Xue Zhong, Jarosław E. Krol, Eva M. Top, and Stephen M. Krone. Accounting for mating pair formation in plasmid population dynamics. Journal of Theoretical Biology, 262(4):711–719, 2010. ISSN 00225193. doi: 10.1016/j.jtbi.2009.10.013.

[36] Jonathan H Bethke, Adam Davidovich, Li Cheng, Allison J Lopatkin, Wenchen Song, Joshua T Thaden, Vance G Fowler, Minfeng Xiao, and Lingchong You. Environmental and genetic determinants of plasmid mobility in pathogenic escherichia coli. Science advances, 6(4):eaax3173, 2020.

[37] Kirsten Riber Philipsen, Lasse Engbo Christiansen, Henrik Hasman, and Henrik Madsen. Modelling conjugation with stochastic differential equations. Journal of theoretical biology, 263(1): 134–142, 2010.

[38] Fang Lei, Yeshi Yin, Yuezhu Wang, Bo Deng, Hongwei David Yu, Lanjuan Li, Charlie Xiang, Shengyue Wang, Baoli Zhu, and Xin Wang. Higher-Level Production of Volatile Fatty Acids In Vitro by Chicken Gut Microbiotas than by Human Gut Microbiotas as Determined by Functional Analyses. Applied and Environmental Microbiology, 78(16):5763–5772, aug 2012. ISSN 0099-2240. doi: 10.1128/AEM.00327-12.

[39] Barry G Hall, Hande Acar, Anna Nandipati, and Miriam Barlow. Growth Rates Made Easy. Molecular Biology and Evolution, 31(1):232–238, jan 2014. ISSN 0737-4038. doi: 10.1093/molbev/mst187.

[40] Jesse B Alderliesten, Sarah J N Duxbury, Mark P Zwart, J Arjan G M de Visser, Arjan Stegeman, and Egil A J Fischer. Effect of donor-recipient relatedness on the plasmid conjugation frequency: a meta-analysis. BMC Microbiology, 20(1):135, dec 2020. ISSN 1471-2180. doi: 10.1186/s12866-020-01825-4. URL https://doi.org/10.1186/s12866-020-01825-4https://bmcmicrobiol.biomedcentral.com/articles/10.1186/s12866-020-01825-4.

[41] Tatyana A. Sysoeva, Youlim Kim, Jonathan Rodriguez, Allison J. Lopatkin, and Lingchong You. Growth-stage-dependent regulation of conjugation. AIChE Journal, 66(3):1–10, mar 2020. ISSN 0001-1541. doi: 10.1002/aic.16848. URL https://onlinelibrary.wiley.com/doi/10.1002/aic.16848.

[42] Paul E Turner. Phenotypic plasticity in bacterial plasmids. Genetics, 167(1):9–20, 2004.

